# Defining host factors required for dengue virus NS1-induced endothelial hyperpermeability and endothelial glycocalyx layer disruption

**DOI:** 10.64898/2026.11.16.766930

**Authors:** Laurentia V. Tjang, Jordan Quellec, Xinyi Feng, Felix Pahmeier, Vasiliya Kril, Kayla Leung, Rémi Hénocque, Samhita Bhat, Jeffrey Li, Pedro H. Carneiro, Scott B. Biering, Eva Harris

## Abstract

Dengue virus (DENV) is a mosquito-borne positive-sense RNA virus belonging to the *Flaviviridae* family that poses a significant threat to global health. Clinical manifestations of DENV infection range from febrile to severe disease characterized by vascular leakage. The secreted DENV nonstructural protein 1 (NS1) contributes to endothelial dysfunction through interactions with endothelial cells, leading to disruption of the endothelial glycocalyx and intercellular junction complexes (IJC). However, host factors required for NS1-triggered endothelial dysfunction are not well understood. It is also unclear if NS1 highjacks different host factors and signaling pathways for glycocalyx and IJC disruption. To address these questions, we used a targeted genetic approach to identify host factors required either for NS1-triggered endothelial glycocalyx layer (EGL) disruption, as assessed by immunofluorescence assays, or for IJC disruption, as measured by transendothelial electrical resistance. After demonstrating the decoupled nature of EGL and IJC disruption in our models, we genetically depleted host factors with known roles in regulating endothelial barrier function and/or involvement in NS1 pathogenesis, including heparanase, cathepsin L, sialidases, and the PI3K/AKT/mTOR and β-catenin signaling pathways. Our results reveal a complex genetic landscape where NS1-induced disruption of EGL and IJCs involves both unique and shared host factors. Further, our data identify a direct role for β-catenin in NS1-induced endothelial dysfunction *in vitro* and *in vivo*. Overall, this work advances understanding of host factors required for DENV NS1-triggered endothelial dysfunction, revealing distinct intracellular pathways that are highjacked by NS1 to modulate barrier function, thus providing new insights into dengue-associated vascular leak.

**Importance:** Dengue virus (DENV) is the most widespread arbovirus in the world, and its severe forms are characterized by endothelial dysfunction and vascular leakage. The DENV non-structural protein 1 (NS1) plays a significant role in disrupting the endothelial glycocalyx (EGL) and mislocalizing intercellular junction complexes (IJCs), two processes essential for endothelial dysfunction. However, the host factors involved in NS1-induced endothelial dysfunction, as well as the overlap between EGL and IJC dysfunction, remain unclear. To address this gap, we genetically disrupted various host factors involved in pathways modulating endothelial barrier function. Our data revealed a complex genetic landscape in which EGL and IJC are mechanistically decoupled and in which NS1-induced disruption of these barriers involves both specific and shared host factors. This work presents a new research framework enabling a better understanding of the molecular mechanisms underlying NS1-induced pathogenesis and, consequently, the identification of new therapeutic approaches for severe forms of dengue.

## Introduction

Dengue is the most prevalent mosquito-borne viral disease in humans and a major public health burden, with an estimated 50-100 million cases occurring annually caused by the four dengue virus serotypes (DENV1-4) (1, 2). Clinical manifestations of dengue range from mild febrile-like illness to severe, potentially life-threatening cases associated with endothelial dysfunction contributing to vascular leak, a hallmark of severe dengue disease (2–4). Despite considerable effort, no specific therapeutics are currently FDA-approved for the treatment of dengue. Three tetravalent vaccines have been licensed in some countries: Dengvaxia^®^ (which has been discontinued), Qdenga^®^ (TAK-003), and Butantan-DV in Brazil (NIH TV-003). However, ongoing concerns about the potential risk of antibody-dependent enhancement (ADE) exemplify the challenges associated with current dengue vaccination strategies targeting the DENV envelope (E)(5). For these reasons, it is crucial to examine other viral targets of importance for DENV pathogenesis, such as viral proteins that contribute to pathogenic manifestations of severe dengue including vascular leak (4, 6–9).

DENV, recently renamed *Orthoflavivirus denguei*, is a small enveloped virus belonging to the *Flaviviridae* family, with a single positive-stranded RNA genome of ∼11kb (3, 10). Upon host cell infection, translation of the DENV RNA genome yields a single polyprotein precursor that undergoes co- and post-translational cleavage by host and viral proteases (11). This process yields distinct viral proteins, including nonstructural protein 1 (NS1), a key virulence factor in flavivirus biology. While intracellular dimeric NS1 is essential for viral replication, the soluble oligomeric form of NS1 is secreted into the bloodstream, where it mediates immune evasion, activates immune cells, and triggers endothelial hyperpermeability and vascular leak (6, 7). Although severe dengue-associated vascular leak was historically attributed to a "cytokine storm" induced by immune cell activation or uncontrolled viral replication, recent evidence highlights direct NS1-mediated endothelial dysfunction as a contributing factor (6, 7, 12). Indeed, soluble DENV NS1 alone is sufficient to induce endothelial barrier breakdown *in vitro* and vascular leak *in vivo* and thus can serve as a pathogenic mediator of severe disease (6, 13, 14). Consistent with this, serum concentrations of NS1 from various flaviviruses correlate with clinical severity (15–17). Mirroring flavivirus tissue tropism, NS1-triggered endothelial dysfunction has been shown to facilitate systemic viral dissemination, allowing the virus to reach permissive cells in its target organs (18–22). Importantly, vaccination against NS1 or passive transfer of anti-NS1 antibodies is protective in mouse models of vascular leak, viral dissemination and lethal DENV challenge, highlighting the central role of NS1 in pathogenesis and viral dissemination (6, 22–24).

Mechanistically, soluble NS1 initiates endothelial barrier dysfunction through interactions with various cell types, including immune cell populations and endothelial cells. NS1 interactions with immune cells are reported to trigger the production of proinflammatory cytokines along with other vasoactive intermediates, contributing to a cytokine storm, but can also mediate protection via inflammasome activation (8, 25). Interactions of NS1 with endothelial cells results in endocytosis of NS1 and the activation of intracellular signaling pathways that ultimately disrupt the key structural complexes maintaining endothelial barrier integrity: the endothelial glycocalyx layer (EGL) and intercellular junctional complexes (IJCs) (26, 27). The EGL and IJC are two distinct cell structures regulating transcellular and paracellular permeability, respectively. The EGL is a dense network of proteins and glycans lining the luminal surface of endothelial cells, composed of glycoproteins bearing acidic oligosaccharides and terminal sialic acid, as well as membrane-bound proteoglycans decorated with glycosaminoglycan side chains, including heparan sulfate (26, 28). Positioned at the apical cell surface, this layer acts as a physical barrier and critical interface regulating mechanotransduction and governing interactions between blood components and the vessel wall (26, 28, 29). Thus, the EGL acts as a key luminal gatekeeper that protects the endothelial barrier by regulating cellular signaling, macromolecule extravasation, and transcellular endothelial transport (26, 30). EGL degradation contributes to endothelial dysfunction by disrupting transcellular permeability and releasing bioactive glycocalyx fragments that further promote inflammation, junctional remodeling, and vascular hyperpermeability (31–33). In addition, the extracellular matrix (ECM), a network of proteins providing structural and biochemical support to cells, promotes endothelial cell maturation and the assembly of IJCs thereby supporting establishment of the endothelial barrier (34, 35). IJCs, consisting of a protein scaffold that includes tight junction (TJ) and adherens junction (AJ) proteins, are intercellular structures that mediate cell-cell adhesion and endothelial barrier function by sealing the endothelium and inhibiting passive paracellular diffusion (27, 36). The maintenance and remodeling of the EGL and IJCs are dynamic processes that regulate many biological responses, including reactions to growth factors, angiogenesis, wound healing, and diverse stress conditions (36, 37). Therefore, the EGL and IJCs play complementary roles protecting against endothelial dysfunction, with the EGL protecting the apical cell surface and IJCs sealing the paracellular space. Accordingly, endothelial dysfunction can be measured by monitoring EGL integrity through staining for glycan components and visualization by immunofluorescence assay (IFA), indirectly reflecting transcellular permeability, and by evaluating IJC integrity via transendothelial electrical resistance (TEER) assays, reflecting paracellular permeability and enabling the assessment of endothelial hyperpermeability (29, 38).

While numerous studies have characterized the functional consequences of NS1 interactions with endothelial cells, knowledge of the host factors and signaling pathways required for NS1-induced barrier dysfunction remains limited (7, 39–41). Our previous studies investigating NS1-mediated barrier dysfunction suggest that NS1 may hijack host cellular pathways, triggering the activation of host factors regulating barrier function (6, 18, 20, 42–46). However, previous studies have relied almost entirely on small-molecule inhibitors rather than genetic approaches, leaving potential off-target effects unaddressed and limiting our understanding of specific host factors in NS1-triggered endothelial dysfunction (39, 42–45). Finally, while NS1 triggers the disruption of both the EGL and IJCs, some studies rely on a single readout, usually TEER, thereby overlooking a critical question in the field, namely, whether the pathways governing EGL and IJC disruption are overlapping or distinct (39, 44, 45).

Following NS1 internalization into endothelial cells, evidence suggests that NS1 triggers upregulation of both protein expression and activity of cathepsin L, heparanase, and neuraminidases (42, 43). The activation of additional enzymes, such as matrix metalloproteinase-9 (MMP-9) and glycogen synthase kinase 3β (GSK-3β), has also been implicated in NS1-triggered barrier dysfunction (20, 44, 46). Furthermore, studies suggest the potential involvement of upstream regulators of cell growth and autophagy during DENV infection, including the PI3K/AKT/mTOR (PAM) signaling axis (47–50). While these host factors seem to be required for the transient disruption of the EGL or IJCs, the interconnection between these NS1-triggered events remains unclear and has not been confirmed genetically. In this study, we genetically dissect three cassettes of pathway- or function-related host factors (comprising 11 gene products in total) known to maintain endothelial barrier integrity, assessing their relative contributions to DENV2 (hereafter referred to as DENV) NS1-mediated endothelial dysfunction using both EGL and TEER readouts. Our work identifies host factors specifically involved in either EGL disruption and/or endothelial hyperpermeability and demonstrates that these processes are mechanistically distinct and can be regulated by different NS1-triggered pathways. Taken together, our study advances knowledge of NS1 pathogenesis by genetically dissecting the distinct roles of numerous host proteins in NS1-mediated endothelial barrier dysfunction, providing a basis for the potential development of host-directed therapeutics to prevent severe forms of dengue disease.

## Results

### EGL and IJC disruption are decoupled events *in vitro*

DENV NS1 interactions with endothelial cells result in at least two events compromising barrier integrity, including EGL and IJC disruption. We sought to investigate if the pathways and host factors that trigger these events are coupled or uncoupled. To address this question, we used human pulmonary microvascular endothelial cells (HPMECs) due to the importance of pulmonary vascular leak in dengue disease severity and the well-established impact of DENV NS1 on HPMEC endothelial function (18, 42, 43, 51, 52). Moreover, the pulmonary vascular glycocalyx is reported to play an important role in maintaining vascular homeostasis (53–55). We dissected the role of DENV NS1 on endothelial dysfunction after CRISPR-Cas9 knockout of candidate host factors by two previously defined readouts (6, 18, 42, 43). First, transcellular integrity was evaluated by quantifying sialic acid residues on glycoproteins and heparan sulfate on proteoglycans, two major components of the EGL, through IFA (37). In parallel, paracellular permeability was monitored by quantifying the integrity of the IJC via a TEER assay **(Fig. 1A)**.

**Figure 1.**
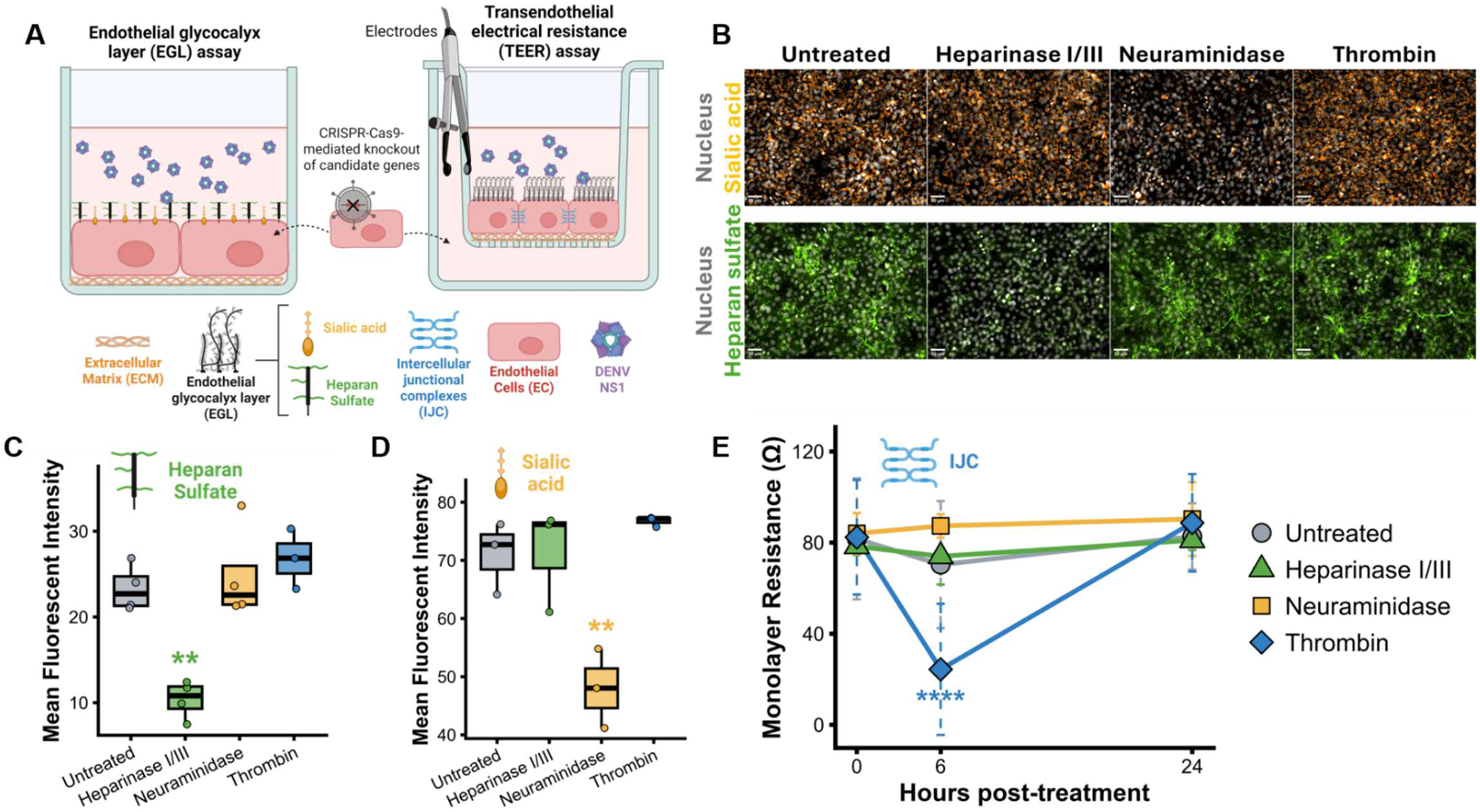
Disruption of the EGL and IJC are decoupled events in HPMECs. **(A)** Cartoon depicting assays measuring endothelial barriers during DENV NS1 treatment. Transcellular permeability was monitored by quantifying EGL glycan components sialic acid and heparan sulfate by IFA quantified as mean fluorescence intensity (MFI) (left). Paracellular permeability was monitored by measuring transendothelial electrical resistance (TEER) of HPMECs seeded in transwell inserts to evaluate the integrity of IJC (right). **(B)** Representative IFA images of HPMECs treated with Heparinase I/III (0.5 U/mL), Neuraminidase (1 U/mL), or thrombin (1 U/mL) for 6 hours as indicated, with sialic acid in yellow (WGA-647), heparan sulfate in green, and nuclei in grey. Images are representative of at least three independent experiments (scale bar, 50µm)**. (C-D)** Quantification of B visualized as MFI. (one-way ANOVA, ** P<0.01). **(E)** TEER measurements on HPMECs with the indicated treatments and time points. Enzyme concentrations as in B. Results representative of three independent experiments are expressed as monolayer resistance (Ω, normalized on blank inserts) and compared to the untreated control for each time point (mean ± 95% confidence interval, one-way ANOVA,**** P<0.0001).

The EGL and IJC are reported to play distinct roles in regulating endothelial biology (26, 27). To confirm this functional distinction in our *in vitro* system, we evaluated if specific disruption of one barrier component impacted the others. To determine if EGL degradation is decoupled from IJC-mediated hyperpermeability measured by TEER, we treated HPMECs with enzymes degrading EGL components: heparinase I/III to cleave heparan sulfate and neuraminidase to remove sialic acid **(Fig. 1B)**. As a positive control for IJC disruption, we used thrombin, a protease known to induce transient endothelial hyperpermeability by triggering signaling pathways that alter IJC localization (56–58). First, we confirmed by IFA that enzymatic treatments induced significant and specific depletion of heparan sulfate and sialic acid **(Fig. 1C-D)**. However, enzymatic degradation of either heparan sulfate or sialic acid had no significant impact on TEER values compared to the untreated control condition **(Fig. 1E)**. Conversely, while thrombin induced a significant and transient drop in monolayer resistance at 6 hours post-treatment, thrombin-mediated IJC disruption did not affect the integrity of either EGL components **(Fig. 1B-D)**. Taken together, these results suggest that *in vitro* disruption of transcellular (EGL) and paracellular (IJC) permeability can be decoupled. This functional separation permits us to study the role of candidate host factors for their role in facilitating EGL and/or IJC disruption *in vitro*.

### Barrier-remodeling enzymes are differentially involved in NS1-triggered EGL disruption and endothelial hyperpermeability

The glycoproteins and proteoglycans that comprise the EGL and the ECM are dynamically modified by enzymes that maintain endothelial integrity and homeostasis (59). Cathepsin L (CTSL), a member of the cathepsin protease family, plays a major role in endothelial homeostasis and exhibits intracellular lysosomal proteolytic activity as well as specific extracellular functions leading to the remodeling of EGL, ECM, or IJC components (26, 60–63). Moreover, CTSL has been specifically shown to mediate the activation of heparanase (HPSE), an endoglycosidase that cleaves heparan sulfate within the EGL and ECM (64–66). Matrix metalloproteinase 9 (MMP-9) belongs to the zinc-metalloproteinase family, capable of degrading EGL, ECM, or IJC components directly (26, 67–70). Pharmacologically, each of these enzymes has been implicated in NS1-mediated endothelial dysfunction, but whether they have a dual role in NS1-triggered EGL and IJC disruption is unclear (42). Given their known biological roles and previous inhibitor studies, we hypothesized that CTSL, HPSE, and MMP9 would differentially contribute to NS1-mediated EGL degradation and IJC disruption.

To test this hypothesis, we utilized CRISPR-CAS9 knockout technology to ablate expression of the genes encoding these proteins in HPMECs. We compared the capacity of the knockout HPMECs to respond to DENV NS1 treatment compared to non-targeting guide (gNT)- transduced control HPMEC lines. We confirmed reduced protein expression (knockout) for CTSL and HSPE in this study, while MMP9 protein knockout performed with the same guides has been previously shown **(Fig. 2A)** (71). We then checked if our knockout HPMECs exhibited an altered endothelial barrier and found no significantly different levels of sialic acid or heparan sulfate determined by IFA nor differences in TEER readings compared to HPMEC/gNT controls, suggesting that knockouts did not impair basal EGL formation or monolayer resistance **(Fig. 2B/D/F)**.

**Figure 2.**
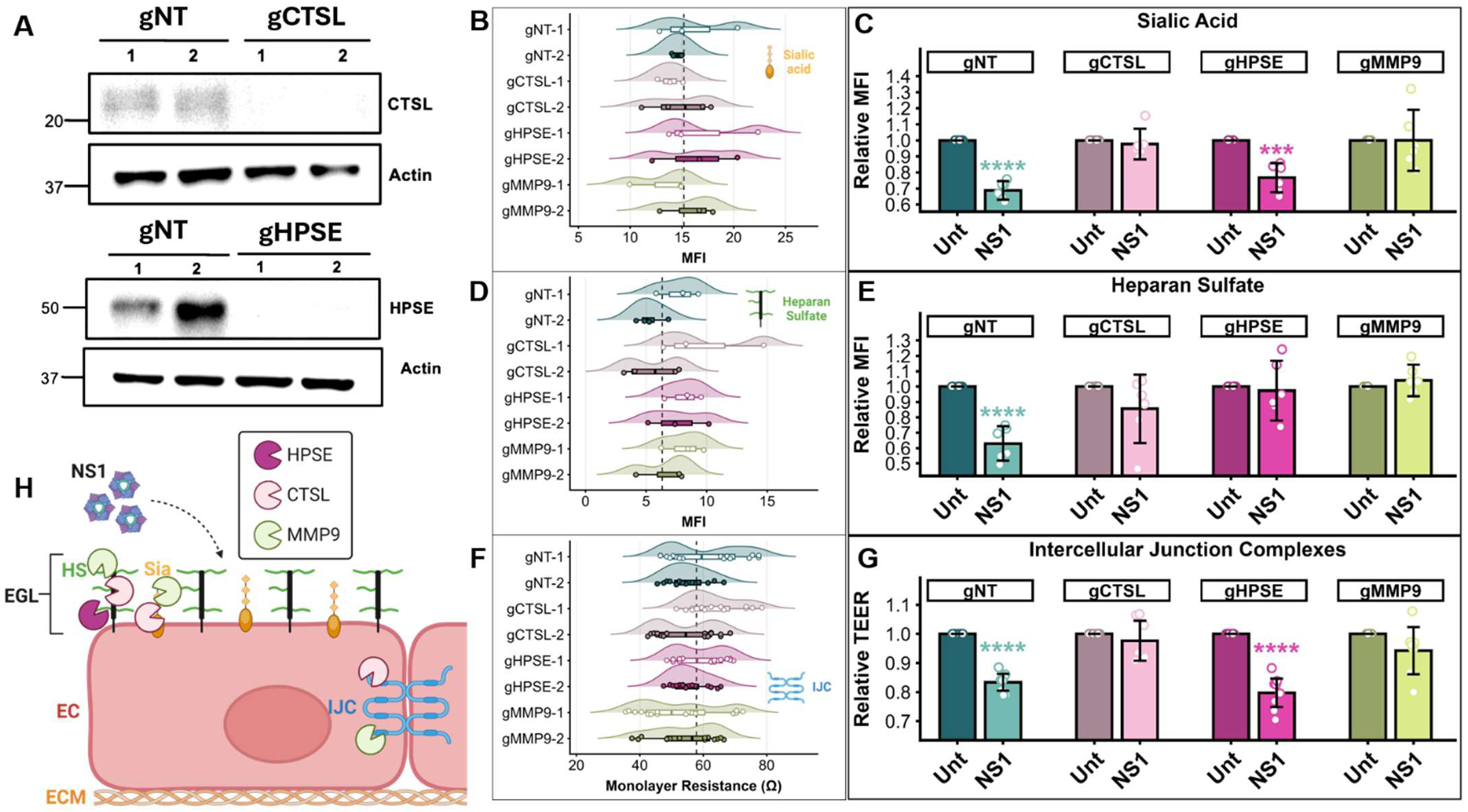
Barrier-modulating enzymes are differentially involved in NS1-triggered endothelial dysfunction. **(A)** Western blot analysis of HPMECs transduced with lentiviruses encoding Cas9 and single guide RNAs targeting genes encoding CTSL (gCTSL) and HPSE (gHPSE), or control non-targeting guides (gNT) as indicated (two independent guides per target). **(B, D and F)** Baseline barrier readings for genetically modified HPMECs (two independent guides, −1 and −2) by measuring sialic acid staining **(B)**, heparan sulfate staining **(D)**, or electrical resistance by TEER **(F)**. Dashed lines indicate the mean barrier values of gNT-1 and gNT-2; one-way ANOVA comparisons were non-significant. **(C, E, and G)** The indicated genetically modified (g) HPMEC were evaluated for EGL integrity (sialic acid in **C** and heparan sulfate in **E**) by immunofluorescence microscopy and for IJC barrier integrity by transendothelial electrical resistance (TEER in **G**) 6 hours post-DENV NS1 treatment (10 µg/mL). Values were normalized to untreated (Unt) conditions for each cell line. Results are from at least three independent experiments merging two independent sgRNA per gene target and expressed as mean ± 95% confidence interval (two-way ANOVA compared to untreated control for each cell line; *** P<0.001, **** P<0.0001). **(H)** Cartoon depicting the roles of barrier-modulating enzymes cathepsin L (CTSL), heparanase (HPSE), and matrix metalloproteinase 9 (MMP9) in NS1-triggered EGL and IJC dysfunction (IJC, Sia or HS). EGL: endothelial glycocalyx layer, HS: heparan sulfate, Sia: sialic acid, EC: endothelial cell, ECM: extracellular matrix.

HPMEC/gNT and knockout cells were then treated with DENV NS1, and the impact on endothelial barriers was assessed by EGL and TEER assays. Compared to untreated conditions for each cell line, HPMEC/gCTSL and HPMEC/gMMP9 knockout HPMECs were no longer sensitive to NS1-mediated sialic acid and heparan sulfate degradation, in contrast to HPMEC/gNT controls, suggesting that both CTSL and MMP9 are required for EGL disruption **(Fig. 2C and 2E)**. Consistent with its enzymatic substrate, HPMEC/gHPSE knockout HPMECs exclusively lost NS1-induced heparan sulfate degradation, highlighting the specificity of HPSE in NS1-induced endothelial dysfunction by selectively driving heparan sulfate cleavage. Further, compared to untreated HPMEC conditions, the TEER assay demonstrated that NS1-mediated hyperpermeability was lost in HPMEC/gCTSL and HPMEC/gMMP9 knockout lines compared to HPMEC/gNT, where NS1-induced hyperpermeability was observed **(Fig. 2G)**. These results confirm the role of CTSL and MMP9 in altering IJCs. However, knocking out HPSE in HPMEC/gHPSE did not prevent NS1-mediated hyperpermeability, suggesting that HPSE is not required for IJC disruption. Taken together, these data indicate that CTSL and MMP9 are required for both NS1-mediated hyperpermeability and EGL degradation, whereas HPSE is specifically restricted to NS1-mediated heparan sulfate cleavage **(Fig 2H)**. These genetic findings demonstrated a broader role for CTSL and MMP9 in NS1-induced endothelial dysfunction while revealing a selective function for HPSE.

### NEU1 and NEU2 are required for NS1-mediated EGL disruption but not for hyperpermeability

Neuraminidases, also known as sialidases, are a family of enzymes that cleave terminal sialic acid residues from glycoproteins in the EGL (66, 72). Neuraminidase 1 (NEU1) is a lysosomal- and plasma membrane-localized sialidase; Neuraminidase 2 (NEU2) is mainly cytosolic; and Neuraminidase 3 (NEU3) is cytoplasmic and membrane-associated (73–75). Our previous work demonstrated that NS1 both triggered sialic acid degradation in addition to increasing expression of NEU1, NEU2, and NEU3 following NS1 treatment in HPMECs (42). This previous study utilized broad-spectrum neuraminidase inhibitors to prevent NS1-mediated barrier dysfunction. However, broad chemical inhibition precluded the identification of specific roles for individual neuraminidases and may have introduced off-target effects that fully inhibited NS1-mediated endothelial dysfunction. Based on their canonical enzymatic functions, we hypothesized that neuraminidases are required for NS1-mediated sialic acid degradation, but not for IJC disruption. Using CRISPR-Cas9, we generated knockout HPMEC lines for each neuraminidase and applied our experimental framework to dissect their individual contributions to NS1-induced endothelial dysfunction.

First, knockout HPMEC lines displayed equivalent baseline MFI (sialic acid, heparan sulfate) and TEER values compared to HPMEC/gNT controls, confirming that gene expression perturbation did not alter basal endothelial biology **(Fig. 3A, 3C, and 3E)**. Upon DENV NS1 treatment, while NS1 induced significant degradation of both sialic acid and heparan sulfate in HPMEC/gNT controls, HPMECs genetically deficient in NEU1 or NEU2 did not exhibit NS1-mediated sialic acid degradation **(Fig. 3B)**. Unexpectedly, HPMEC/gNEU1 and HPMEC/gNEU2 knockout lines were also refractory to NS1-mediated heparan sulfate degradation, indicating that NEU1 and NEU2 are broadly required for overall EGL disruption **(Fig. 3D)**. However, consistent with our initial hypothesis, all neuraminidase-deficient cell lines remained sensitive to NS1-induced hyperpermeability as measured by TEER **(Fig. 3F)**. In contrast, HPMEC/gNEU3 behaved similarly to HPMEC/gNT controls, remaining susceptible to both NS1-mediated hyperpermeability and EGL degradation **(Fig. 3B, 3D, and 3F)**, suggesting that NEU3 is dispensable for NS1-induced endothelial dysfunction. These data collectively suggest that NEU1 and NEU2 are essential for NS1-mediated EGL disruption, affecting both sialic acid and heparan sulfate disruption, while NEU3 seems non-essential for NS1 activity. Importantly, these data suggest that these neuraminidases are not involved in IJC disruption and endothelial hyperpermeability **(Fig. 3G)**.

**Figure 3.**
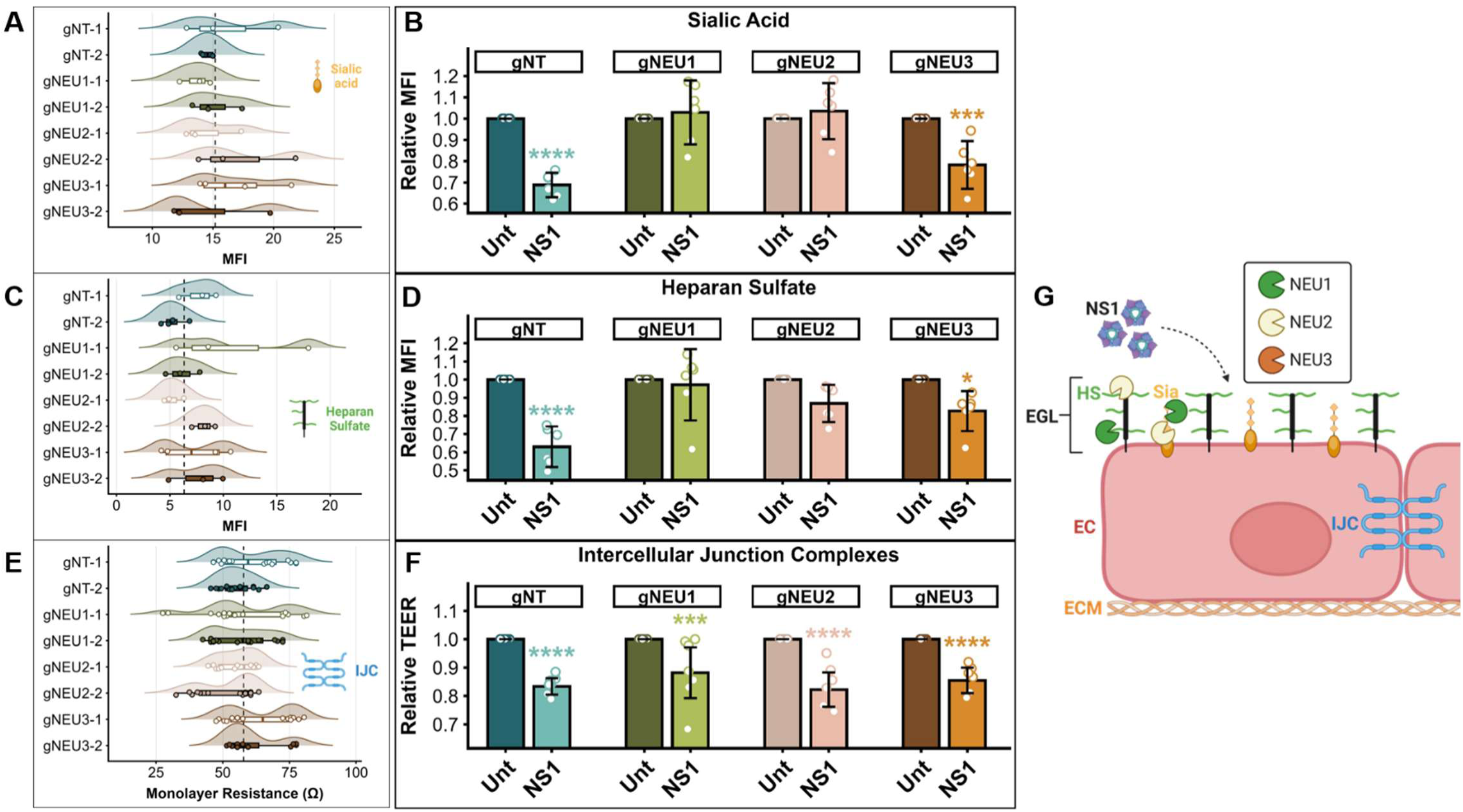
NEU1 and NEU2 are required for NS1-mediated EGL disruption but not for paracellular permeability. (A, C, and E) Baseline barriers of the indicated genetically modified (g) HPMECs (two independent single guide RNAs (sgRNA), −1 and −2) with mean fluorescence intensity values of sialic acid **(A)** and heparan sulfate **(C)** measuring EGL disruption and TEER values **(E)** measuring paracellular barrier integrity, as indicated. Dashed lines indicate the mean of gNT-1 and gNT-2; non-significant by one-way ANOVA. **(B, D, and F)** The indicated genetically modified (g) HPMECs were evaluated for EGL integrity (sialic acid in **B** and heparan sulfate in **D**) by immunofluorescence microscopy and for IJC barrier integrity by TEER (**F**) 6 hours post-DENV NS1 treatment (10 µg/mL). Values were normalized to untreated (Unt) conditions for each cell line. Results are from at least three independent experiments merging two independent sgRNAs per gene target and expressed as mean ± 95% confidence interval. Two-way ANOVA compared to untreated control were performed for each cell line (* P<0.05, *** P<0.001, **** P<0.0001). **(G)** Cartoon illustrating the roles of neuraminidases in NS1-triggered endothelial dysfunction (IJC, Sia or HS). EGL: endothelial glycocalyx layer, HS: heparan sulfate, Sia: sialic acid, EC: endothelial cell, ECM: extracellular matrix.

### PI3K, AKT, and mTOR are required specifically for NS1-triggered EGL disruption

The PAM signaling axis is a major intracellular pathway regulating cell cycle, survival, motility, proliferation, and autophagy (76). Phosphatidyl-inositol 3 kinases (PI3Ks) are a family of plasma membrane-associated lipid kinases, AKT is a serine/threonine kinase, and mammalian-target of rapamycin (mTOR) is a serine/threonine protein kinase belonging to the PI3K-related protein kinase (PIKK) family (77–79). Previous studies using inhibitors and siRNA silencing demonstrated that autophagy induction in endothelial cells, potentially mediated through Macrophage Migration Inhibitory Factor (MIF) secretion, is required for NS1-induced endothelial dysfunction (47). These findings implicate the potential involvement of the PAM signaling pathway in NS1 activity, leading us to hypothesize its requirement for both forms of NS1-induced endothelial dysfunction.

To test the role of PAM components in NS1-mediated endothelial dysfunction, we used two independent sgRNAs per gene to knock out *PIK3R1* (encoding the regulatory p85 subunit crucial for PI3K activity), *AKT1*, and *MTOR* (77). Protein depletion was confirmed by Western blot **(Fig. 4A)**, and the functional effects of these gene deletions on baseline endothelial biology were evaluated relative to the gNT control **(Fig. 4B, 4D, and 4F)**. None of the genetically modified HPMEC lines exhibited a significant reduction in basal MFI for cell surface sialic acid and heparan sulfate compared to HPMEC/gNT, indicating that gene deletion did not impair baseline EGL formation **(Fig. 4B and 4D)**. Interestingly, the HPMEC/gMTOR-1 line (sgRNA 1, mTOR-1) exhibited an increase in baseline monolayer resistance compared to the HPMEC/gNT control **(Fig. 4F)**. However, because this difference was modest ∼10 Ω and NS1 still induced a measurable drop in relative TEER in both HPMEC/gMTOR lines, confirming a consistent phenotypic response, both sgRNAs were retained in the analysis **(Fig. 4G)**.

**Figure 4.**
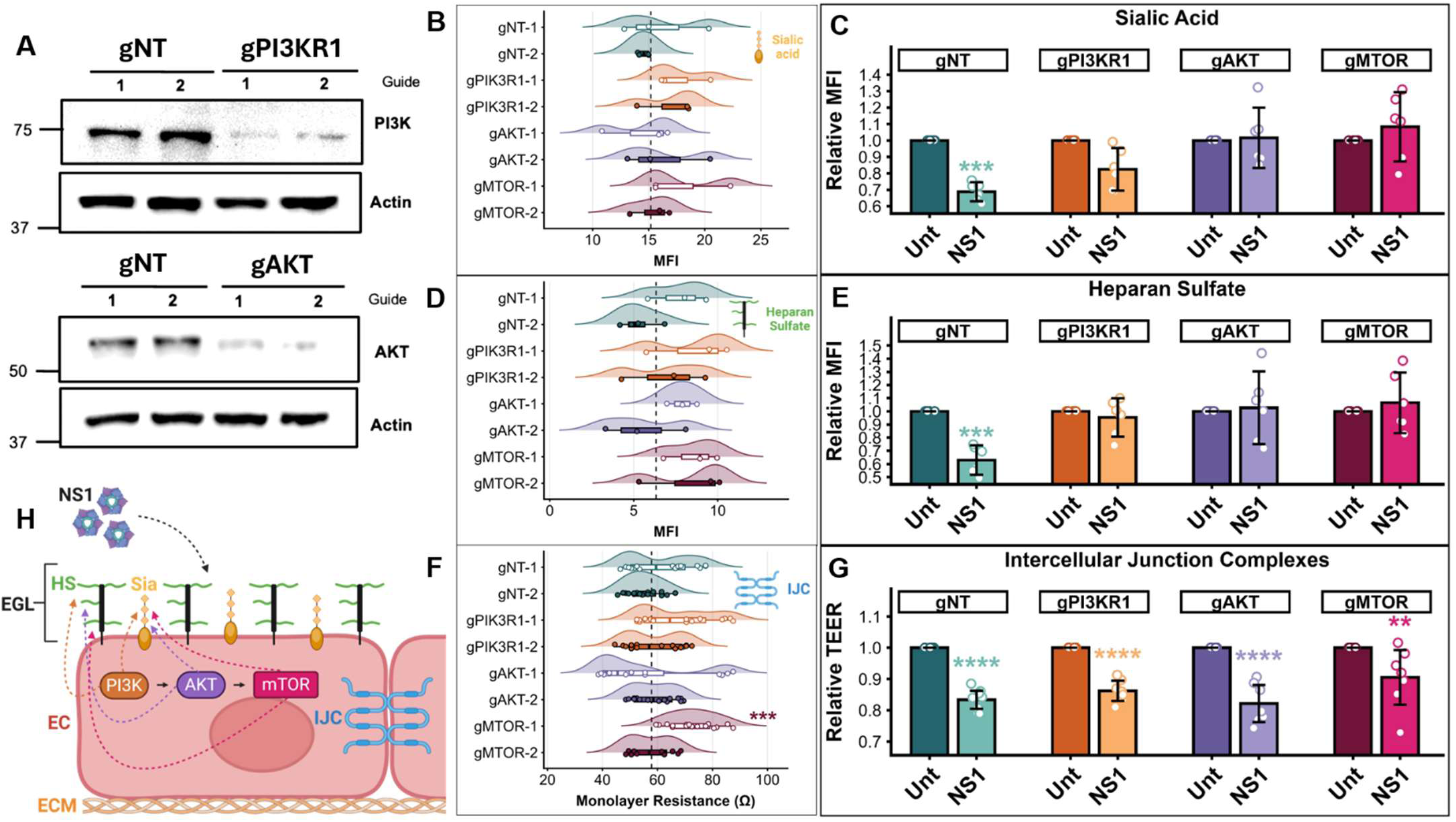
The PI3K-AKT-mTOR (PAM) components are required for NS1-mediated EGL disruption but not endothelial hyperpermeability. **(A)** Western blot analysis of HPMECs transduced with lentiviruses encoding Cas9 and guide RNAs targeting genes encoding PI3KR1 (gPI3KR1) and AKT1 (gAKT) or control non-targeting guides (gNT) as indicated (two independent guides per target). **(B, D, and F)** Baseline barriers of the indicated genetically modified (g) HPMECs (two independent single guide RNAs (sgRNA), −1 and −2) with mean fluorescence intensity values of sialic acid **(B)** and heparan sulfate **(D)** measuring EGL disruption and TEER values **(F)** measuring paracellular barrier integrity, as indicated. Dashed lines indicate the mean of gNT-1 and gNT-2; non-significant by one-way ANOVA. **(C, E, and G)** The indicated genetically modified HPMECs were evaluated for EGL integrity (sialic acid in **C** and heparan sulfate in **E**) by immunofluorescence microscopy and for IJC barrier integrity by TEER (**G**) 6 hours post-DENV NS1 treatment (10 µg/mL). Values were normalized to untreated (Unt) conditions for each cell line. Results are from at least three independent experiments merging two independent sgRNAs per gene target and expressed as mean ± 95% confidence interval. Two-way ANOVA compared to untreated control was performed for each cell line (** P<0.01, *** P<0.001, **** P<0.0001). **(H)** Cartoon illustrating the roles of PAM pathway proteins in NS1-triggered endothelial dysfunction (IJC, Sia or HS). EGL: endothelial glycocalyx layer, HS: heparan sulfate, Sia: sialic acid, EC: endothelial cell, ECM: extracellular matrix.

First, we evaluated sialic acid and heparan sulfate levels relative to untreated conditions for each cell line treated with NS1. In contrast to HPMEC/gNT controls, where NS1 treatment triggered significant degradation of both sialic acid and heparan sulfate, none of the knockout lines (*PIK3R1*, *AKT1*, or *MTOR*) exhibited significant EGL degradation following NS1 exposure **(Fig. 4C and 4E).** Interestingly, all knockout cell lines remained fully sensitive to NS1-induced hyperpermeability as measured by relative TEER drop **(Fig. 4G)**, indicating that PAM pathway components are required for EGL disruption but not for NS1-mediated IJC disruption and hyperpermeability. These data suggest that the PAM signaling pathway selectively regulates a distinct aspect of NS1-induced endothelial dysfunction **(Fig 4H)**.

### GSK3β is involved in NS1-mediated endothelial hyperpermeability, whereas β-catenin is involved in overall NS1-mediated barrier dysfunction

Our previous work highlighted the involvement of GSK3β in *in vitro* and *in vivo* endothelial dysfunction induced by DENV NS1 in umbilical vein and brain microvascular endothelial cells (44). GSK3β is a serine/threonine kinase that plays a central role in modulating the phosphorylation and function of β-catenin (80). β-Catenin is a dual-function protein whose subcellular localization and phosphorylation status dictate its specific role, including regulation of the endothelial barrier (81). Indeed, non-phosphorylated β-catenin acts as a structural component of AJ within the IJC complex or translocates to the nucleus for transcriptional regulation, whereas phosphorylated β-catenin is targeted for cytoplasmic degradation (82). Our previous work using chemical inhibitors showed that GSK3β is required for NS1-mediated hyperpermeability, hypothesized to be via the mislocalization of β-catenin (44). Given the central role of GSK3β in regulating β-catenin, alongside the key function of β-catenin in IJC stability, we hypothesized that both GSK3β and β-catenin are essential for NS1-mediated hyperpermeability, but dispensable for EGL degradation.

To test this hypothesis genetically, we utilized CRISPR-Cas9 to generate GSK3β and β-catenin knockout HPMECs using two independent sgRNAs per gene. After confirming protein depletion by Western blot **(Fig. 5A)** and verifying that knockouts did not impair basal endothelial biology compared to HPMEC/gNT controls **(Fig. 5B, 5D, and 5F)**, we conducted EGL and TEER assays following NS1 treatment. As predicted, HMPEC/gGSK3β behaved similarly in the EGL assay to HPMEC/gNT controls, showing that NS1 treatment exhibited significant degradation of both sialic acid and heparan sulfate relative to untreated conditions **(Fig. 5C and 5E)**. Unexpectedly, β-catenin knockout HPMECs did not exhibit NS1-mediated EGL degradation compared to untreated conditions, as sialic acid and heparan sulfate fluorescence intensity were not significantly affected by NS1 treatment. In the TEER assay, both GSK3β and β-catenin knockout HPMECs showed abolished NS1-mediated hyperpermeability compared to HPMEC/gNT controls, where NS1 treatment retained significant hyperpermeability relative to untreated conditions **(Fig. 5G)**. These results support our hypothesis that GSK3β and β-catenin are required for NS1-mediated IJC disruption and subsequent paracellular hyperpermeability.

**Figure 5.**
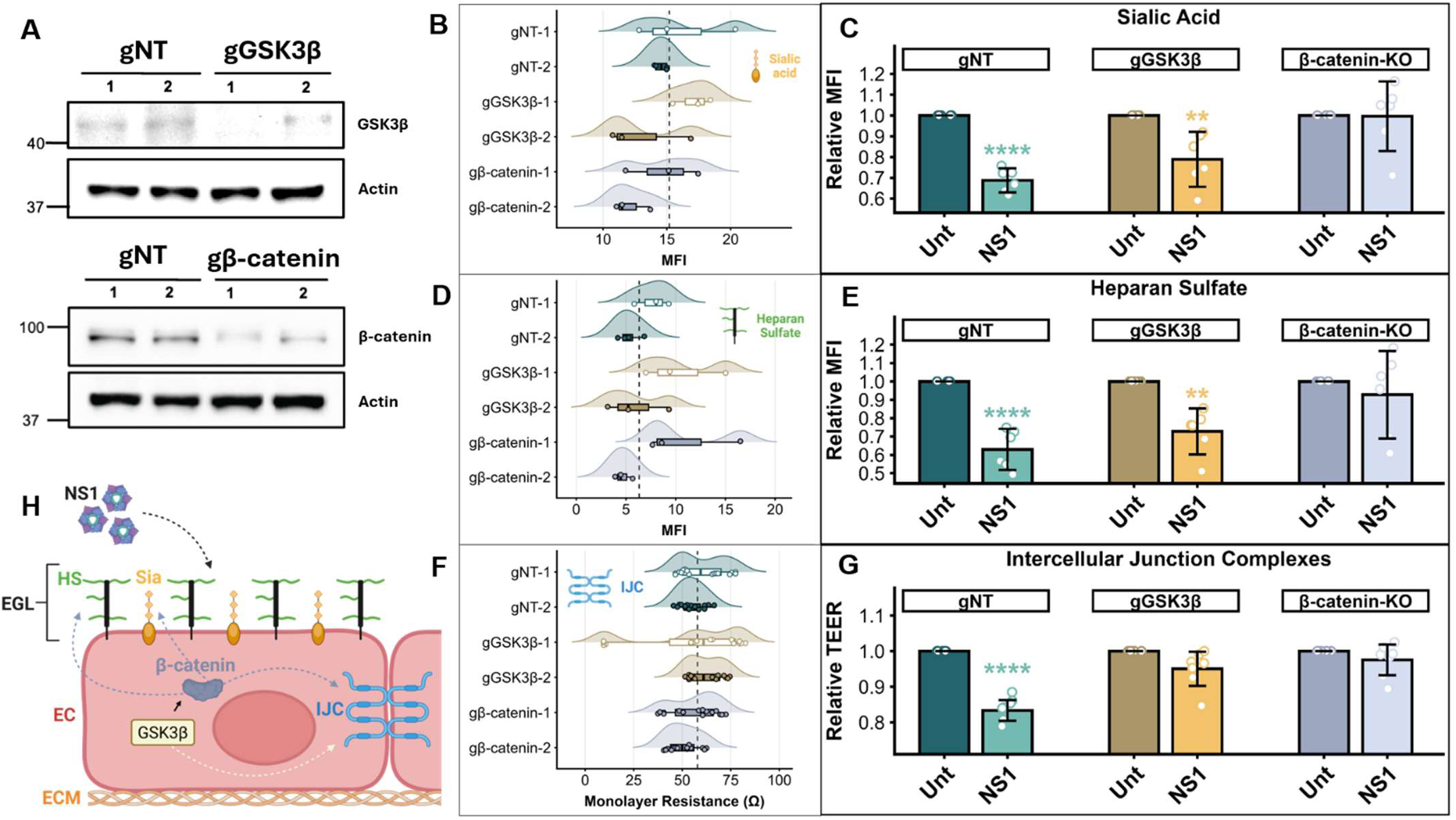
GSK3β is required only for NS1-triggered barrier hyperpermeability, and β-catenin is required for broad endothelial dysfunction. (A) Western blot analysis of HPMECs transduced with lentiviruses encoding Cas9 and guide RNAs targeting genes encoding GSK3β (gGSK3β) and β-catenin (g β-catenin), or control non-targeting guides (gNT) as indicated (two independent guides per target). (B, D, and F) Baseline barriers of the indicated genetically modified (g) HPMECs (two independent single guide RNAs (sgRNA), −1 and −2) with mean fluorescence intensity values of sialic acid (B) and heparan sulfate (D) measuring EGL disruption and TEER values (F) measuring paracellular barrier integrity, as indicated. Dashed lines indicate the mean of gNT-1 and gNT-2; non-significant by one-way ANOVA. (C, E, and G) The indicated genetically modified HPMECs were evaluated for EGL integrity (sialic acid in C and heparan sulfate in E) by immunofluorescence microscopy and IJC barrier integrity by TEER (G) 6 hours post-DENV NS1 treatment (10 µg/mL). Values were normalized to untreated (Unt) conditions for each cell line. Results are from at least three independent experiments merging two independent sgRNAs per gene target and expressed as mean ± 95% confidence interval. Two-way ANOVA compared to untreated control was performed for each cell line (** P<0.01, **** P<0.0001). (H) Cartoon illustrating the roles of signaling proteins GSK3β and β-catenin in NS1-triggered endothelial dysfunction (IJC, Sia or HS). EGL: endothelial glycocalyx layer, HS: heparan sulfate, Sia: sialic acid, EC: endothelial cell, ECM: extracellular matrix.

Altogether, these data revealed a surprisingly broader role for β-catenin in overall NS1-mediated barrier dysfunction, participating in both EGL degradation and endothelial hyperpermeability. Furthermore, these genetic findings confirmed and refined our previous observations that GSK3β is required for NS1-induced endothelial dysfunction, revealing its specific restriction to induction of hyperpermeability **(Fig. 5H)**.

### Small-molecule inhibition of β-catenin prevents NS1-mediated vascular leak *in vivo*

Following up on our observation that β-catenin is a key driver of NS1-triggered endothelial dysfunction, we tested its role in NS1-triggered vascular leak *in vivo*, where endothelial barrier conditions are more physiologically complex. We first validated the *in vitro* efficacy of the chemical inhibitor iCRT14, which inhibits β-catenin responsive transcription **(Fig. 6A)**. In wild-type parental HPMECs, pre-treatment with iCRT14 prevented both NS1-mediated sialic acid degradation and hyperpermeability, whereas vehicle pre-treatment (DMSO) failed to block either phenotype. Thus, this small-molecule inhibitor recapitulated the phenotypic outcomes observed in our genetic knockout models for β-catenin.

**Figure 6.**
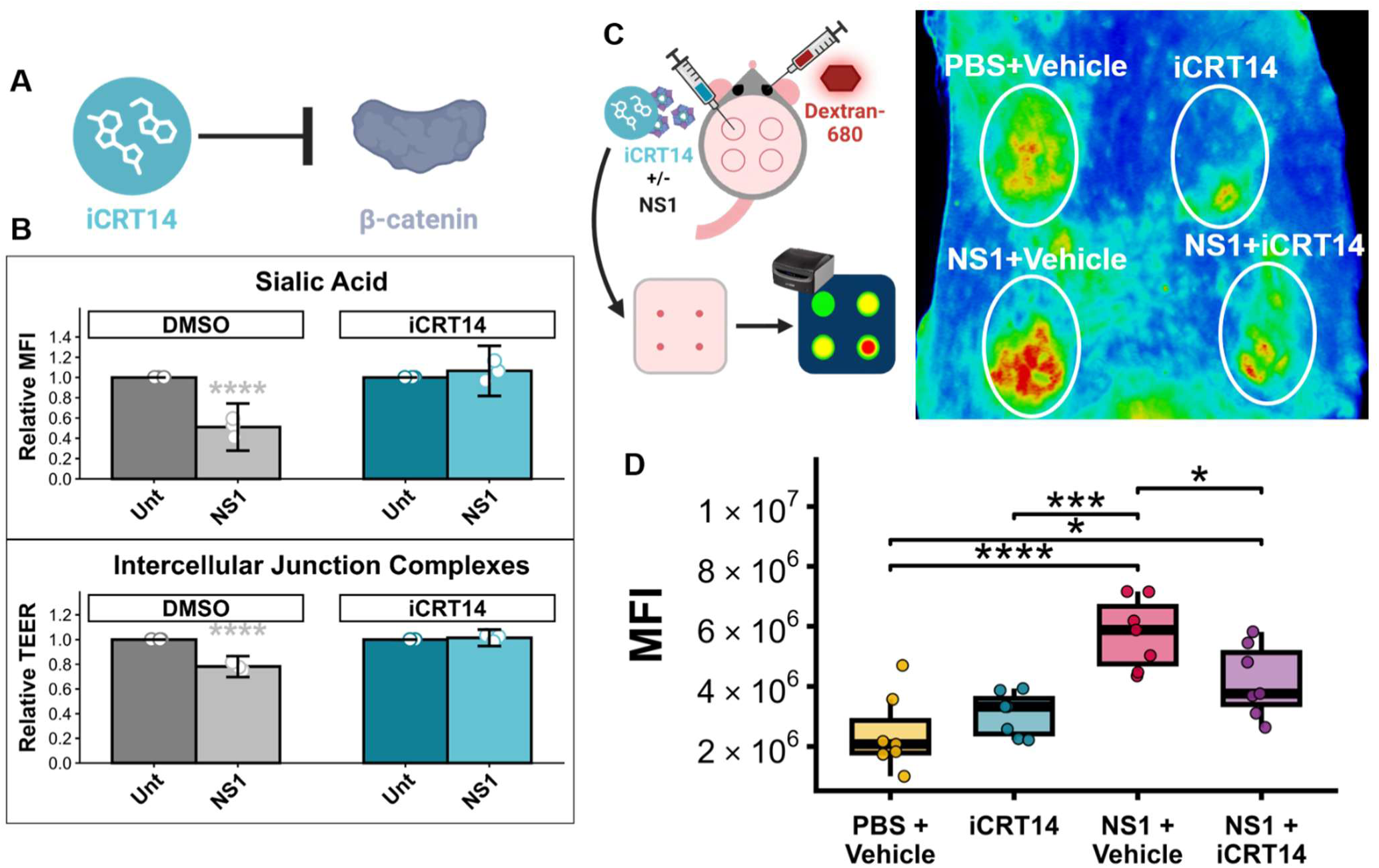
β-Catenin-targeting small molecule iCRT14 inhibits NS1-mediated endothelial dysfunction *in vitro* and vascular leak *in vivo*. (A) Schematic diagram showing chemical inhibition of β-catenin by iCRT14. (B) EGL disruption and endothelial hyperpermeability assays of HPMECs treated with 2 µM of iCRT14 or DMSO as a vehicle control 30 minutes before treatment with DENV NS1 (10 µg/mL) followed by assessment of sialic acid levels by IFA (top) or electrical resistance by TEER (bottom) 6 hours post-treatment. Top is expressed as relative MFI normalized to untreated (Unt) conditions (mean ± 95% confidence interval, two-way ANOVA compared to untreated for each treatment; **** P<0.0001). Bottom is expressed as relative TEER normalized to untreated conditions (mean ± 95% confidence interval, two-way ANOVA compared to untreated for each treatment; **** P<0.0001). (C) Murine intradermal vascular leak assay with the indicated treatments (15 µg of DENV NS1 and 1 µM of iCRT14) administered intradermally into the shaved dermises of mice followed by intravenous administration of the tracer 10-kDa Alexa Fluor 680-conjugated dextran. 2 hours post-administration, dermises were collected and fluorescence was assessed. (D) Quantification of B expressed as mean fluorescence intensity (MFI) (one-way ANOVA, * P<0.05, *** P<0.001, **** P<0.0001).

To test the contribution of β-catenin in mice *in vivo*, we utilized our localized dermal vascular leak (BackID) assay to quantify NS1-induced vascular leak (23, 43). Briefly, intradermal injections of DENV NS1 in the presence or absence of inhibitor were performed on the dorsal dermis of wild-type C57BL/6J mice. Local fluid extravasation, serving as a proxy for vascular leak, was quantified using dermal accumulation of fluorescent signal from the tracer dye 10-kDa Dextran conjugated to Alexa fluor 680, delivered intravenously prior to intradermal challenge **(Fig. 6B)**. As expected, β-catenin inhibition by iCRT14 treatment partially abrogated NS1-induced endothelial dysfunction and subsequent vascular leakage *in vivo* **(Fig. 6B and C)**. Quantification demonstrated that intradermal injection of PBS or iCRT14 alone did not induce significantly different fluid extravasation, whereas co-injection of NS1 with iCRT14 significantly reduced vascular leakage compared to NS1 co-injected with vehicle control, while maintaining a level significantly higher than the PBS baseline **(Fig. 6C)**. These results suggest that transcriptional inhibition of β-catenin partially restricts NS1-mediated endothelial barrier disruption *in vivo*. Taken together, these *in vivo* findings indicate that β-catenin plays a key role in NS1-mediated vascular leak.

## Discussion

Our study reveals differential involvement of numerous host factors in DENV NS1-mediated EGL degradation and/or hyperpermeability. First, we confirmed *in vitro* the current hypothesis that EGL-associated transcellular permeability and IJC-related paracellular permeability constitute two independent barriers that regulate endothelial biology in the model used in this study. Moreover, using a genetic approach, we found that MMP9, CTSL, and β-catenin play a central role in overall NS1-mediated barrier dysfunction, including both EGL and IJC disruption. However, some host factors are involved in either EGL disruption or endothelial hyperpermeability. For instance, we found that the PAM pathway plays an important role only in NS1-mediated EGL disruption but not endothelial hyperpermeability. Conversely, genetic ablation demonstrated that GSK3β is only involved in NS1-mediated hyperpermeability but not EGL disruption. Interestingly, among neuraminidases, only NEU1 and NEU2, but not NEU3, are critical for NS1-mediated EGL degradation (both sialic acid and heparan sulfate) in HPMECs, while HPSE was observed to be required only for heparan sulfate disruption (**Fig. 7A and 7B)**.

**Figure 7.**
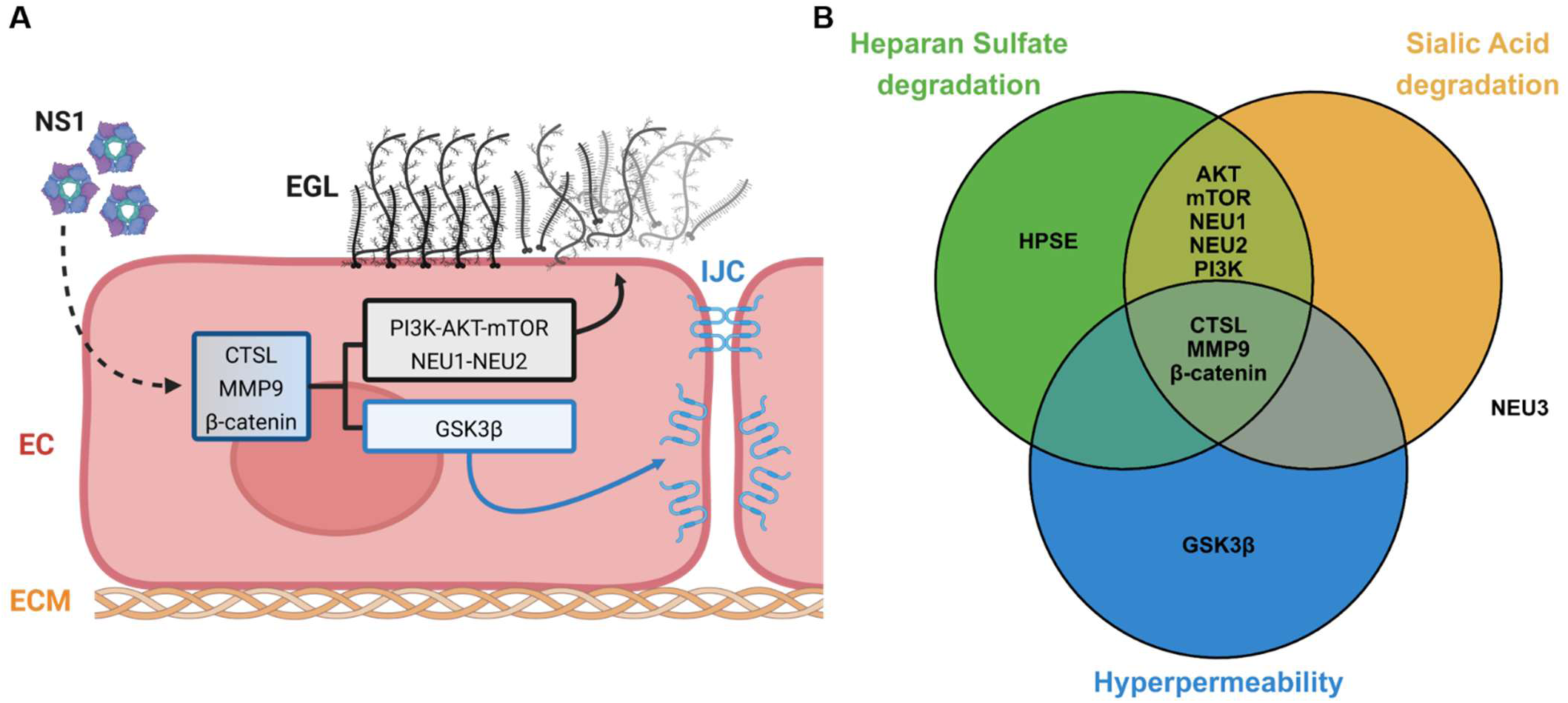
Summary illustration of the overlapping and distinct roles of host factors in NS1-mediated EGL and IJC disruption. **(A)** Host factors are grouped into three categories exhibiting genetic involvement individually in EGL or IJC dysfunction or playing a role in both NS1-triggered processes. **(B)** Summary of results visualized as a Venn-diagram, with NEU3 exhibiting no role in the tested readouts.

The PAM pathway is a highly conserved signaling cascade in eukaryotic cells that regulates the cell lifecycle (83). Interestingly, PAM is known to regulate endothelial cell EGL biology, typically serving a protective rather than a disruptive role to support endothelial function (26). Nevertheless, it has been shown that the PAM pathway can regulate EGL degradation as well as activation of MMP9, which is known to drive EGL degradation (26, 84, 85). Thus, the involvement of PAM in NS1 pathogenesis remains concordant with its canonical signaling functions. Because our data suggest PAM is specific to NS1-mediated EGL disruption while MMP9 is also linked to IJC destabilization, it is possible that PAM functions independently of MMP9 in the context of NS1 or that additional modulating factors are required, such as NF-κB, to fully explain this complexity. DENV NS1 has shown to increase NF-κB activation in human liver cells (HepG2), supporting this hypothesis (86). DENV NS1 has also been demonstrated to induce MMP9 expression in endothelial cells through activation of the NF-κB pathway and to interact directly with MMP9, promoting its recruitment to the IJC complex (β-catenin, Zonula Occludens [ZO] proteins) and triggering its destabilization (46). This latter result supports the idea that the PAM and NF-κB pathways can be activated by DENV NS1 in endothelial cells. Interestingly, even in a PAM knockout context where activation of the NF-κB transcription factor and upregulation of MMP9 expression are restricted, NS1 still induces hyperpermeability. This suggests that NS1 interaction with other host factors is sufficient to induce endothelial hyperpermeability, whereas the PAM pathway is necessary for EGL degradation.

This work also highlights the dual functions in transcellular and paracellular permeability of the endopeptidases CTSL and MMP9, which are known to disrupt components of both the EGL and ECM, the latter regulating IJC scaffolding. In contrast, we defined a precise function for HPSE in NS1-mediated endothelial dysfunction, specifically in EGL degradation where, concordant with its endoglycosidase function, HPSE exclusively mediates heparan sulfate degradation. Interestingly, a recent study in humans also showed that infection with the Puumala Hantavirus induces EGL degradation characterized by specific HPSE-mediated heparan sulfate degradation alongside broader activity of MMP9 on EGL component degradation through syndecan-1 cleavage (68). Extracellular targets of MMP9 and CTSL are primarily known to be associated with the extracellular matrix (ECM), such as gelatin, collagen, or fibronectin, whereas direct degradation of the EGL by these proteases is less well-characterized (60, 61, 70, 87, 88). Interestingly, syndecan ectodomains are cleaved by MMPs during EGL shedding, releasing various EGL components associated with these proteoglycans, including heparan sulfate and chondroitin sulfate (87, 89). For MMP9, only syndecan ectodomain shedding has been demonstrated as a direct contribution to EGL breakdown (70, 87). Similarly, aside from its indirect role in EGL degradation through the processing and activation of HPSE, CTSL is not known to directly digest structural proteins of the EGL (61, 65). Thus, for both proteases, the degradation of sialic acid residues on EGL glycoproteins is likely not a direct enzymatic effect, but rather an indirect effect on another cell signaling cascade, making it important to study the broader involvement of these enzymes in EGL modulation. Indeed, both proteases have been reported to digest various glycoproteins, and it would be important to determine whether the cleavage activity of MMP9 and CTSL acts directly on extracellular substrates or indirectly via the activation of neuraminidases or other EGL-degrading processes, such as reactive oxygen species (ROS) production, thereby explaining the wider EGL degradation observed in our study (90–92). In support of this indirect mechanism, NEU1 activation in immune cells has been shown to depend on MMP9 function, providing a rationale for an indirect role of MMP9 in sialic acid degradation (93). Similarly, CTSL is essential for the maturation of CTSA, a core component of the multienzyme complex that regulates NEU1 activity (93–96).

Further, our study highlights activity of NEU1 and NEU2 in NS1-mediated EGL degradation, whereas NEU3 did not appear to be involved. Four mammalian sialidases have been described (NEU1–4), which differ by tissue expression pattern, subcellular localization, and functional role (97–100). NEU3 and NEU4 are primarily known to process gangliosides at the neural cell surface, showing high expression in skeletal muscle and liver but low expression in the lung. This low physiological expression in pulmonary tissue as well as ganglioside-specific activity likely explains why NEU3 is dispensable for NS1-mediated EGL degradation in HPMECs, a pattern that may also extend to NEU4 and that requires further exploration (97–100). NEU1 is the most abundant and ubiquitously expressed neuraminidase, acting primarily within lysosomal and plasma membrane compartments. In contrast, NEU2, which acts mainly in the cytosol but is also associated with the plasma membrane, is expressed at lower levels and remains less understood. Nonetheless, both sialidases process cell-surface glycoproteins and glycopeptides (101–104). These functional properties corroborate our experimental findings demonstrating the involvement of NEU1 and NEU2 in the observed sialic acid degradation of EGL. Interestingly, and concordant with our earlier hypothesis, NEU2 expression has been shown to be regulated by the PAM pathway, providing a potential link between NEU2 activity and PAM-mediated regulation of NS1-induced EGL degradation (105). Moreover, our previous work highlighted upregulation of NEU1, NEU2, and NEU3 expression in HPMECs following DENV NS1 treatment, reinforcing these observations (42). Here, upregulation of NEU3 expression alone does not appear to dictate its involvement in EGL degradation in HPMECs, perhaps related to its specific preference for ganglioside substrates (97–100).

However, similarly to MMP9 and CTSL, involvement of sialic acid-degrading enzymes NEU1 and NEU2 in heparan sulfate disruption is surprising. Because NEU1, NEU2, CTSL, and MMP9 all appear to be required for both NS1-mediated sialic acid and heparan sulfate breakdown, we postulate that their actions are synergistic, complementary, and organized into an enzymatic activation cascade leading to enhanced EGL destabilization. This notion of enzymatic synergy is supported by comparing exogenous sialidase treatment with DENV NS1 challenge: while exogenous neuraminidase specifically removed sialic acid without affecting heparan sulfate integrity, NEU1 or NEU2 knockout during NS1 treatment prevented the degradation of both components. This suggests that NS1-triggered EGL destruction requires concerted cross-activation between neuraminidases, proteases (MMP9, CTSL), and endoglycosidases (HPSE), where disrupting a single player halts overall EGL breakdown. However, the precise hierarchy of this cooperative cascade remains to be fully clarified.

This work also helps to refine the role of GSK3β as a key signaling factor leading specifically to endothelial hyperpermeability, whereas unexpectedly, its canonical partner, β-catenin, is involved in overall NS1-mediated endothelial dysfunction. We originally hypothesized that β-catenin would be important only for maintaining IJC integrity, given our previous observation that DENV NS1 triggers the mislocalization of β-catenin from the IJC (44). Indeed, cell membrane-associated β-catenin binds to cadherins within adherens junctions, directly supporting barrier function (106). However, β-catenin may not act solely through its direct structural role at adherens junctions during NS1-mediated disruption, given that the β-catenin knockout cell line maintained basal paracellular permeability comparable to gNT controls. In canonical Wnt signaling, GSK3β forms part of the destruction complex that mediates the phosphorylation and subsequent proteasomal degradation of cytoplasmic β-catenin (107, 108). Given the functional interplay between GSK3β and β-catenin within the Wnt pathway, we hypothesize that Wnt signaling plays a central role in NS1-mediated endothelial dysfunction, as supported by our *in vitro* and *in vivo* findings. GSK3β restricts the transcriptional activity of β-catenin by limiting its cytoplasmic concentration (107, 108). When GSK3β is inhibited or absent, non-phosphorylated β-catenin functions as a transcription factor, driving the expression of target genes involved in diverse cellular processes. Although this transcriptional activity is essential for blood-brain barrier maturation and maintenance via claudin regulation, several downstream targets (such as VEGF, IL-6, and TNFα) are known to induce transient endothelial hyperpermeability and EGL degradation, potentially through secondary NF-κB activation, though this additional mechanism is speculative and needs to be clarified (107–110). Taken together, these data suggest that β-catenin acts as a transcription factor driving the upregulation of genes associated with transient endothelial dysfunction and breakdown of both the EGL and IJC.

An important aspect of this study is the *in vitro* functional independence demonstrated between transcellular and paracellular permeability, which enabled us to dissect the precise contributions of distinct host factors to NS1-mediated endothelial dysfunction. Our data indicate that, in the context of DENV NS1, EGL and IJC disruption are distinct processes but share overlapping genetic requirements. Although each permeability axis tested in our study was evaluated using a single primary method (TEER or IFA), the specificity of these approaches was validated by targeted degradation of endothelial components (sialic acid degradation by neuraminidase, heparan sulfate degradation by heparinase I/III) and targeted perturbation of IJC by thrombin) -- without functional overlap. However, to strengthen the robustness of our transcellular permeability in future studies, macromolecular diffusion assays and TEER-based evaluations of IJC integrity paired with junctional imaging and passive paracellular flux assays could be used.

However, the independence of EGL and IJC disruption in general endothelial dysfunction and vascular leak *in vivo* is not clear. Indeed, some studies report that syndecan-1, a product of EGL shedding, is positively correlated with vascular leak, whereas other authors describe a negative correlation or no correlation between them (66, 68, 111–115). Other components of the EGL, such as sialic acid, are also believed to enhance systemic vascular leak (26, 43, 113, 116). Additionally, EGL degradation weakens the endothelial barrier by removing apical protection, thereby exposing the cell surface to blood components that potentially contribute to overall endothelial dysfunction, notably inflammatory components such as immune cells or cytokines (26, 113). Several factors can explain the independence of these two permeability pathways, and more importantly, the discrepancy between *in vitro* and *in vivo* observations, notably differences in the kinetics of endothelial dysfunction and EGL maturation. Indeed, in comparison to the hyperpermeability resulting from the early and direct impact of NS1 on endothelial cells, we hypothesize that EGL shedding gradually increases the systemic and bystander impact of NS1 on the vasculature, resulting in a delayed and amplified vascular leak by weakening the endothelial transcellular barrier (26, 42, 113). Furthermore, it has been shown that endothelial cells *in vitro* harbor an immature and thinner EGL compared to cells *in vivo* (117). We can then hypothesize that immature EGL degradation *in vitro* is not sufficient to induce the full destabilization of endothelial function described *in vivo*, even if the main components are present. Understanding how different host factors coordinate barrier dysfunction and their kinetics in NS1-triggered vascular leak *in vivo* is undoubtedly complex and represents a critical next step for the flavivirus NS1 field.

Furthermore, as noted previously, the host factors identified in this study pertain specifically to DENV NS1 treatment of pulmonary endothelial cells. Given the distinct tissue expression patterns of host factors implicated in NS1-mediated endothelial dysfunction, such as NEU3 and NEU4, which exhibit higher activity in the brain and liver, it is reasonable to hypothesize that intracellular signaling pathways may vary across organotypic endothelial cells and potentially among NS1 flaviviral origins that exhibit different tissue tropism (18, 97–100). Nevertheless, certain central factors could remain shared across cell types and viral species, as demonstrated with GSK3β in umbilical cells with ZIKV NS1 and in brain endothelial cells with DENV and ZIKV NS1 (18, 44). Interestingly, different flavivirus NS1 proteins (DENV, ZIKV, JEV, TBEV) and endothelial cell types (brain, dermal, umbilical, pulmonary) have been shown to engage additional pathways and host factors that contribute to overall NS1-mediated endothelial dysfunction, including VE-cadherin and RhoA phosphorylation, miR-145-5p and type I IFN signaling, the p38-MAPK pathway, the TNFα pathway, TLR4, and reactive oxygen species (39, 45, 118–120). To our knowledge, this current study represents the first precise genetic mapping of host factors to distinct permeability routes during NS1-mediated endothelial dysfunction. However, further investigation is needed to establish both organ- and virus-specific host mechanisms alongside the core pathways common to the NS1-induced phenotype.

In conclusion, our study sheds light on the differential pathways and host factor dependencies utilized by DENV NS1 to trigger endothelial barrier dysfunction. Moreover, our results are complementary to previous findings, allowing us to clarify the specific functions of enzymes previously described in NS1-mediated endothelial dysfunction. Our genetic dissection approach, rather than relying on chemical inhibitors that potentially elicit off-target effects, permits us to understand more granularity in the roles of each host factor (42, 43). Most importantly, these findings provide valuable leads for dissecting intracellular signaling cascades and identifying key host factors involved in specific aspects of NS1-mediated pathogenesis. This process is inherently complex, engaging multiple cellular components that contribute to distinct aspects of endothelial permeability, as well as exhibiting potential organ and viral specificities. Overall, this work provides fundamental insights into the host factor requirements of DENV NS1-mediated pathogenesis, shedding light on the mechanisms which could facilitate NS1-mediated pathogenesis in DENV infection, and thus leading to vascular leak in severe form the disease as well as promoting viral dissemination and its consequent pathology.

## Materials and Methods

### Cell lines and proteins

293T cells were obtained from ATCC and maintained in DMEM, high glucose, GlutaMAX™ (Gibco) supplemented with 10% FBS (Corning) and 1% penicillin/streptomycin (Gibco) (D10 medium) at 37°C with 5% CO_2_. Human pulmonary microvascular endothelial cells (HPMECs) (line HpMEC-ST1.6R) were kindly provided by Dr. J.C. Kirkpatrick at Johannes Gutenberg University, Germany (121). HPMECs were cultured in endothelial basal growth medium 2 supplemented with an endothelial growth medium 2 supplemental bullet kit (EGM2, Lonza). Cells were maintained at 37°C with 5% CO_2_. Recombinant NS1 protein from DENV-2 (Thailand strain 16681) produced in HEK293 cells was purchased from Native Antigen Co. (United Kingdom).

### CRISPR gRNAs design, lentivirus production and HPMEC transduction

The CRISPR-Cas9 KO protocol was adapted from Zhang et al., using lentiCRISPRv2 lentivirus construct (Addgene plasmid #52961; http://n2t.net/addgene:52961; RRID:Addgene_52961)(122). Two distinct guide RNAs per gene of interest were selected from the Brunello CRISPR knockout guide library and cloned into lentiCRISPRv2 plasmid using a Golden Gate Assembly method **(Supplemental Table 1)** (123). For second-generation lentivirus production, vectors encoding gRNAs and Cas9 (lentiCRISPRv2) were transfected into 293T cells using a polyethyleneimine (PEI) transfection protocol along with the lentivirus packaging vector psPAX2 and the pseudotyping vector pMD2.G. Lentivirus was then collected at 24, 36, and 48 hours post-transfection and added to parental HPMECs. HPMECs were incubated for 24-48 hours, then selected during 3 passages in EGM2 medium containing 2 µg/mL of puromycin. To validate depletion of the targeted protein, Western blot was performed by using the following antibodies: mouse anti-Cathepsin L (Thermo Fisher Scientific), rabbit anti-Heparanase 1 (Abcam), rabbit anti-PI3K p85 alpha (Abcam), rabbit anti-AKT (Abcam), rabbit anti-GSK3β (Cell Signaling), mouse anti-β-catenin (Thermo Fisher Scientific), mouse anti-β-actin HRP-conjugated (Santa Cruz Biotechnologies), goat anti-mouse HRP-conjugated (Biolegend), and donkey anti-rabbit HRP-conjugated (Biolegend). Cell lysates were run in SDS-PAGE gel and proteins were probed with primary antibodies diluted 1:1000 and then with HRP-conjugated secondary antibodies diluted at 1:5000. Membranes were then imaged on a ChemiDoc system with Image Lab software v6.1.0. When possible, in absence of sufficiently specific antibodies or a noticeable phenotype, genetic indels were confirmed by barcode sequencing **(Supplemental Table 2, Supplemental Figure 1).**

### Endothelial Glycocalyx Layer (EGL) disruption assay

HPMECs (1×10^5^ cells) were seeded onto gelatin-coated coverslips in 24-well plates for 3 days allowing formation of a confluent monolayer and maturation of the EGL before treatment with soluble NS1 and controls. DENV2 NS1 was added to cells at 10 μg/mL, Heparanase I and III Blend from *Flavobacterium heparinum* (Sigma) at 0.5 U/mL, neuraminidase (sialidase) from *Clostridium perfringens* (Sigma) at 1 U/mL, and human thrombin at 1 U/mL, followed by incubation for 6 hours at 37°C. Cells were fixed with 4% PFA and stained for components of the EGL: sialic acid (Wheat Germ Agglutinin [WGA]- conjugated to Alex Fluor 647) and antibody anti-heparan sulfate (Amsbio 370255-s). WGA-647 (10 µg/mL) was added to live cell medium 1 hour pre-fixation. Heparan sulfate was stained on fixed cells with primary antibody incubation overnight followed by secondary goat anti-mouse IgM conjugated to Alexa 488 (Thermo Fisher Scientific A-21042) (XXX) incubation for 2 hours at RT. Nuclei were stained with Hoechst 33342 (Immunochemistry). Coverslips were mounted onto microscope slides on a drop of ProLong Gold (Thermo Fisher Scientific). Confocal imaging was performed using a Zeiss LSM 710 confocal microscope at the CRL Molecular Imaging Center at UC Berkeley. Fluorescence was quantified as measuring Mean Fluorescence Intensity (MFI) for each field analyzed. Results were expressed as relative MFI by normalizing every condition to the untreated control of the corresponding experiment (MFI sample/MFI untreated).

### Trans-endothelial Electrical Resistance (TEER) Assay

HPMECs (6×10^4^ cells) were seeded into the apical chambers of 24-well transwell polycarbonate membrane inserts (Transwell permeable support, 0.4 µM, 6.5 mm insert; Corning). Medium was changed daily from both the apical and basolateral chambers until a confluent monolayer reached maximum electrical monolayer resistance across the transwell insert. Treatments were directly added to the apical chamber of the transwells in cell medium for 6 hours at the concentrations described above. Electrical resistance values were measured in ohms (Ω) at the indicated time-points using an Epithelial Volt Ohm Meter (EVOM) with “chopstick” electrodes (World Precision Instruments). Untreated cells, as well as transwell inserts containing medium only, were used as negative controls to calculate baseline electrical resistance. Monolayer resistance (Ω) was determined by normalizing TEER values of sample with that of transwells with medium only (Ω_sample_ - Ω_medium_). Relative TEER was calculated as a ratio of resistance values as (Ω_sample_- Ω_medium_)/(Ω_untreated_-Ω_medium_).

### Murine Dermal Vascular Leak Assay

The back intradermal (ID) injection assay was adapted from the Miles assay to assess local vascular hyperpermeability (23, 43). Experiments were performed using six-week-old wildtype female C57BL/6J mice obtained from Jackson Laboratory. Three to 7 days prior to experiments, the dorsal fur of the mice was removed using a hair clipper followed by depilatory cream (Nair, Church & Dwight). On the day of experiment, Alexa Fluor 680-conjugated 10-kDa dextran (1 mg/mL; Sigma) was administered intravenously through retro-orbital injection as a tracer and mice were euthanized with isoflurane 2 hours post-injection. Treatment and control (DMSO diluted in PBS) were intradermally injected into distinct spots on the depilated back. Four back ID injections were performed on each mouse, and the dorsal dermises of mice were excised and flattened on a transparent plastic surface. Local vascular leak was visualized using a LI-COR Odyssey CLx Imaging System, and fluorescence signal was quantified using Image Studio software (LI-COR Biosciences).

### Chemical Inhibitors

The β-catenin transcriptional inhibitor (iCRT14, Tocris) was used in EGL disruption, TEER, and Back ID assays. iCRT14 was used at a concentration of 2 μM for *in vitro* assays and at 1 μM in Back ID experiments. Inhibitors were added 30 minutes prior to NS1 treatment. None of the inhibitor concentrations used exhibited significant cytotoxicity.

### Statistics and graphical representation

All data were plotted and analyzed using RStudio (2025.05.1 Build 513). Experiments were performed in at least 3 biological replicates. For CRISPR-KO comparison, data from the two sgRNAs were merged after verification of consistent endothelial phenotypes between sgRNAs and validation of protein depletion or genetic knockout validation. Each cassette was compared with the same two non-targeting guide (gNT) controls (pooled data), and the corresponding gNT data are displayed across multiple panels. For immunofluorescence microscopy experiments, images of random fields were captured. For all experiments with quantitative analyses, data are displayed as mean ± 95% confidence interval (CI). Statistical significance was evaluated using one-way or two-way ANOVA or the Kruskal-Wallis test for non-parametric data. P-values are indicated only where significant (*p < 0.05, **p < 0.01, ***p < 0.001, and ****p < 0.0001).

## Acknowledgements

We thank Chi Zhu (UC Berkeley) for helpful discussion on experimental design. This work was supported by grant R01 AI168003 (EH) from the National Institute of Allergy and Infectious Diseases of the National Institutes of Health. SBB was supported in part by the Life Science Research Foundation, Coefficient Giving, the Keck Foundation, and the Hartwell Foundation. We would like to thank the CRL Molecular Imaging Center for the confocal imaging experiments conducted there, with support from the Gordon and Betty Moore Foundation. Figures were created with BioRender.com.

## Author contributions

L.V.T., S.B.B., and E.H. conceived the study. L.V.T and S.B.B. designed experiments. L.V.T., X.F., F.P., V.K., R.H., K.L., S.B., P.H.C., and J.L. conducted experiments. L.V.T. and J.Q. performed data analysis. L.V.T., J. Q., S.B.B., and E.H. wrote the original draft of the manuscript and all authors participated in reviewing the manuscript.

## Conflict of interest statement

No potential conflict of interest was reported by the author(s).

## Ethics statement

All mice used in this study were bred in-house in compliance with Federal and University regulations. All animal experiments were approved by the Animal Care and Use Committee (ACUC) of the University of California, Berkeley.

## Data access statement

All data pertinent to this study can be found within the paper, and raw data is available upon request. Standard data and material transfer agreements govern all materials and data used in this study.

